# Genome manipulation by guide-directed Argonaute cleavage

**DOI:** 10.1101/2021.11.17.469050

**Authors:** Shan Huang, Kaihang Wang, Stephen L. Mayo

## Abstract

Many prokaryotic argonautes (pAgos) mediate DNA interference by using small DNA guides to cleave target DNA. A recent study shows that CbAgo, a pAgo from *Clostridium butyricum*, induces DNA interference between homologous sequences and generates double-stranded breaks (DSBs) in target DNAs. This mechanism enables the host to defend against invading DNAs such as plasmids and viruses. However, whether such a CbAgo-mediated DNA cleavage is mutagenic remains unexplored. Here we demonstrate that CbAgo, directed by plasmid-encoded guide sequences, can cleave genome target sites and induce chromosome recombination between downstream homologous sequences in *Escherichia coli*. The recombination rate correlates well with pAgo DNA cleavage activity and the mechanistic study suggests the recombination involves DSBs and RecBCD processing. In RecA-deficient *E. coli* strain, guide-directed CbAgo cleavage on chromosomes severely impairs cell growth, which can be utilized as counter-selection to assist Lambda-Red recombineering. These findings demonstrate the guide-directed cleavage of pAgo on the host genome is mutagenic and can lead to different outcomes according to the function of the host DNA repair machinery. We anticipate this novel DNA-guided interference to be useful in broader genetic manipulation. Our study also provides an in vivo assay to characterize or engineer pAgo DNA cleavage activity.

## INTRODUCTION

Prokaryotic argonaute proteins (pAgos) constitute a diverse protein family (10, 11). Unlike their eukaryotic counterparts, which use small RNA guides to interfere with RNA targets in regulation and defense (12, 13), many pAgos were reported to cleave DNA targets using small guide DNAs (gDNAs) in vitro (1-8). In vivo, several pAgos were shown to protect bacteria from foreign DNAs (14-18), but the defense mechanism, especially whether double-stranded DNA breaks (DSBs) are involved, remained elusive until recently. In-depth insights into the mechanism of pAgo-mediated defense were gained by analyzing CbAgo, a pAgo nuclease from a mesophilic bacterium *Clostridium butyricum*, in *Escherichia coli* as its expression host (9). In that study, a DNA interference pathway was revealed in CbAgo-mediated protection against invading DNAs. First, CbAgo generates and binds gDNAs from plasmids or other multicopy genetic elements. Next, gDNA-bound CbAgo introduces DSBs at the homologous sites, including chromosomal regions, and causes DNA degradation in collaboration with E. coli exonuclease RecBCD. Invader DNAs such as plasmids and phages can thus be targeted and eliminated efficiently through this mechanism.

It has been postulated that pAgo may have potential applications in genome manipulation ever since the discovery of its DNA nuclease activity, which could come with greater flexibility over CRISPR-based methods because pAgo-targetable sites are not limited by the presence of a protospacer adjacent motif (PAM) (1, 19). However, to the best of our knowledge, mutations induced by guide-directed cleavage of pAgos in the host genome have never been firmly established. The observation that CbAgo can be directed to generate DSBs in chromosomes by plasmid-encoded guide sequences (GSs) motivated us to leverage such a mechanism to manipulate *E. coli* genomes. Here we demonstrate that guide-directed CbAgo cleavage can directly induce chromosome recombination between direct repeat sequences, or assist Lambda-Red recombineering in *E. coli* as a counter-selection. The recombination system described here can also serve as an efficient in vivo assay to report or engineer pAgo DNA nuclease activity as we find the recombination rates from different pAgos correlate well with their reported in vitro cleavage activity. These findings demonstrate the potential of establishing a DNA-directed genome editing system using pAgo.

## MATERIAL AND METHODS

### Culture conditions

*Escherichia coli*, cultured in Luria-Bertani (LB) medium and agar, was incubated at 37°C or 30°C. When appropriate, antibiotics were added to the medium at the following final concentrations: ampicillin, 100 μg/ml; chloramphenicol, 20 μg/ml; kanamycin, 35 μg/ml. Bacterial cell growth was monitored periodically by measuring the optical density of culture aliquots at 600 nm.

### Strains and plasmids

*Escherichia coli* strains used in this study are listed in Supplementary Table S1. Plasmids used in this study are listed in Supplementary Tables S2. Oligonucleotides used in this study are listed in Supplementary Tables S3. Procedures for the construction of strains and plasmids are described in Supplementary information.

### Determination of recombination frequency

Cells were transformed with appropriate plasmids and plated on LB plates supplemented with ampicillin. The next day, 5 ml of LB medium supplemented with ampicillin was inoculated with single colony and aerated at 37 °C until OD600 = 0.3-0.4. The temperature was then adjusted to 18 °C and after 30 min protein expression was induced by adding anhydrotetracycline to 200 ng/ml for 16 h. Cultures were then cooled down on ice for 10 min, washed with ice-cold PBS (pH 7.2), resuspended in 5 ml of LB medium supplemented with ampicillin, and recovered at 37 °C for 5 h. Serial dilutions of cells were plated on the LB plates supplemented with appropriate antibiotics to determine cfu.

### Fluctuation analysis

Cells were transformed with appropriate plasmids and plated on LB plates supplemented with ampicillin. The next day, 1 ml of LB medium supplemented with ampicillin and 200 ng/ml anhydrotetracycline was inoculated with single colony and aerated at 37 °C for 12 h before making serial dilutions of cultures and plating on the LB plates supplemented with appropriate antibiotics to determine cfu.

For plasmid-free strain 3×ChikanS, 3×ChikanS_ pal246_ΔsbcCD, and 3×ChikanS_pal246, 1 ml of LB medium was inoculated with single colony and aerated at 37 °C for 12 h before making serial dilutions of cultures and plating on the LB plates without antibiotic or supplemented with kanamycin to determine cfu.

The Ma-Sandri-Sarkar Maximum Likelihood Estimator (MSS-MLE) Method or the Lea-Coulson Method of the Median in the Fluctuation AnaLysis CalculatOR (FALCOR) (25) were used to calculate recombination rates and 95% confidence intervals. The online FALCOR tool is available at https://lianglab.brocku.ca/FALCOR/.

### Flow cytometry analysis

Strain SMR6669 cells were transformed with appropriate plasmids and plated on LB plates supplemented with ampicillin. The next day, 1 ml of LB medium supplemented with ampicillin and 200 ng/ml anhydrotetracycline was inoculated with single colony and aerated at 37 °C for 12 h. Cultures were then washed with ice-cold PBS (pH 7.2), diluted 1:500 into ice-cold PBS (pH 7.2), passed through 40 μm cell strainers, added 1 µg/ml propidium iodide to determine cell viability, and analyzed on a Beckman Coulter Cytoflex S Flow Cytometer. For each experiment, 10^5^ cells per culture and 3 cultures per strain were analyzed.

Flow cytometry data were analyzed using FlowJo software version 10.8.1. To comparatively quantify green cells, a green “gate” was set arbitrarily as the window in which ∼0.9% of the control strain, SMR6669/pEmpty fall, according to the spontaneous SOS induction level (27).

### Lambda-Red recombineering

The kanamycin-resistance cassette was amplified from genomic DNA of SIJ488_ΔlacZ via colony PCR with primers Lambda.Red.F/Lambda.Red.R. Resulting PCR product was gel-purified as dsDNA donor. To calculate mutation efficiency for Lambda-Red recombineering (referred to as the standard recombineering procedure), 5 ml of LB medium was inoculated with single colony of strain SIJ488_ΔrecA and aerated at 37 °C until OD600 = 0.3-0.4. The Lambda-Red genes were then induced with 15 mM L-arabinose for 45 min. The culture was used to prepare electrocompetent cells by washing twice with 10% glycerol and resuspending in 50 μl 10% glycerol. 2 μl mixture of ∼300 ng dsDNA was added to the cells, which were then subject to electroporation and allowed to recover in 1 ml LB for 2 h at 37 °C. Serial dilutions of cells were plated on the LB plates with no antibiotic or supplemented with kanamycin to determine cfu.

To calculate mutation efficiency for CbAgo-assisted Lambda-Red recombineering, cells of strain SIJ488_ΔrecA were transformed with appropriate plasmids and plated on LB plates supplemented with ampicillin. The next day, 5 ml of LB medium supplemented with ampicillin was inoculated with single colony and aerated at 37 °C until OD600 = 0.3-0.4. The Lambda-Red genes were then induced with 15 mM L-arabinose for 45 min. The culture was used to prepare electrocompetent cells by washing twice with 10% glycerol and resuspending in 50 μl 10% glycerol. 2 μl mixture of ∼300 ng dsDNA was added to the cells, which were then subject to electroporation and allowed to recover in 1 ml LB supplemented with ampicillin for 2 h at 37 °C. The recovered cells were diluted into 5 ml LB supplemented with ampicillin and 0.2% glucose and continued growing for 2 hours at 37 °C. The temperature was then adjusted to 18 °C and after 30 min protein expression was induced by adding anhydrotetracycline to 200 ng/ml for 16 h. Upon induction, cultures were cooled down on ice for 10 min, washed with ice-cold PBS (pH 7.2), resuspended in 5 ml of LB medium supplemented with ampicillin, and recovered at 37 °C for 5 h. Serial dilutions of cells were plated on the LB plates supplemented with appropriate antibiotics to determine cfu.

### Statistical analyses

GraphPad Prism 9 was used to evaluate statistical significance. Student’s t-test (two-tailed) was used for the statistical analysis of experiments. *P* values < 0.05 were considered significant.

## RESULTS

### Creation of the recombination system

Previous observation that guide-directed CbAgo cleavage at *E. coli* chromosomes efficiently triggers RecBCD activity (9) inspired us to hypothesize that RecBCD-dependent chromosome recombination should be triggered by guide-directed CbAgo cleavage as well. It has been shown that DSBs introduced by SbcCD cleavage at a 246-bp chromosomal palindrome (pal246) stimulate RecBCD-dependent recombination between two downstream direct repeat sequences (20). To determine whether similar recombination can be induced by guide-directed CbAgo cleavage (Figure 1A, Supplementary Figure S1), we integrated a recombination cassette at cynX locus on *E. coli* strain DL1777, which is ∼6 kb away from the target lacZ locus (Figure 1B). This recombination cassette contains an EM7 promoter and a kanamycin resistance gene whose function is abolished by the insertion of a stop codon array and a 270-bp direct repeat sequence. A recombination event between the two direct repeat sequences removes the insertion, restores the gene function, and confers kanamycin resistance to the host. Considering the introduction of the Chi site, an 8-base 5’-GCTGGTGG-3’ motif recognized by RecBCD (21-24), has been shown to stimulate RecBCD recombination (20), we also incorporated a varied number of Chi sites into the genome, with their 5’ ends oriented towards the recombination cassette (Figure 1B).

**Figure 1.**
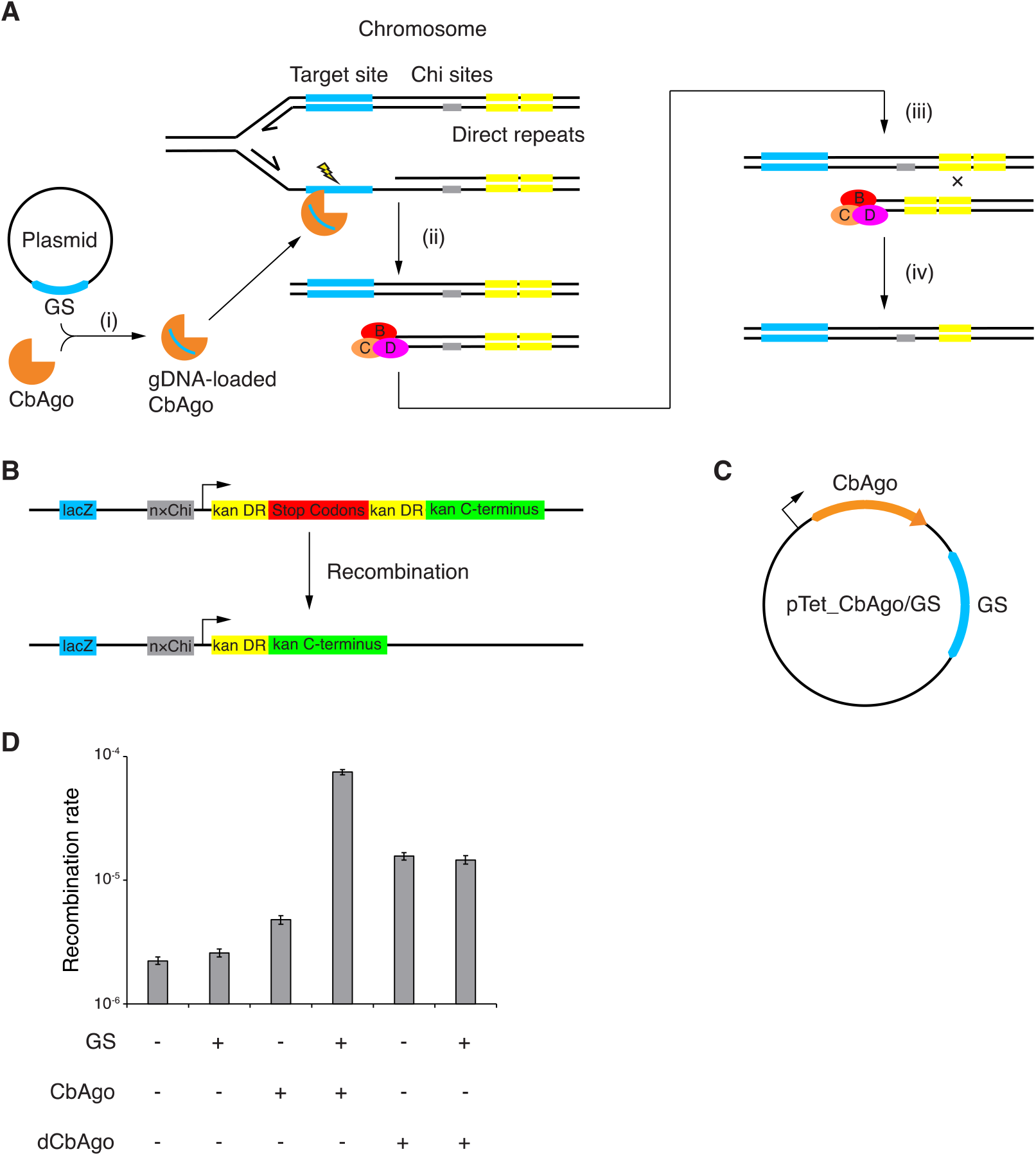
Guide-directed CbAgo cleavage stimulates chromosome recombination. (A) Proposed mechanism for guide-directed CbAgo cleavage and RecBCD-dependent recombination. (i) CbAgo acquires gDNA from plasmid-encoded GS. See Supplementary Figure S1 for the proposed model of gDNA biogenesis. (ii) Guide-directed CbAgo cleavage of lagging strand during chromosome replication generates DSB and (iii) triggers RecBCD binding and processing of chromosome DNA. (iv) Recognition of Chi sites by RecBCD attenuates its DNA degradation activity and facilitates recombination between sister chromosomes via RecA. (B) Structure of the recombination cassette. See materials and methods for DNA sequences. kan, kanamycin resistance gene. DR, direct repeat. (C) Structure of the CbAgo targeting plasmid. (D) Recombination rates of strain 3×ChikanS in different genetic contexts determined by fluctuation analysis from eight independent cultures. Error bars represent 95% confidence intervals.

To target the lacZ locus, we created a targeting plasmid pTet_CbAgo/GS encoding a CbAgo expression cassette under the control of a tetracycline-inducible promoter (pTet), and a 1000-bp GS homologous to lacZ gene (Figure 1C). Importantly, GS is the only sequence on the plasmid (except for an 80-bp rrnB T1 terminator sequence) that is homologous to the genome, assuring only the lacZ locus will be effectively targeted. For controls, plasmids with no GS, no CbAgo gene, or neither, were created. To determine the dependence of CbAgo cleavage activity, we created plasmids encoding a CbAgo mutant (dCbAgo: CbAgo D541A-D611A) that contains mutations of two catalytic residues in its active site which were previously shown to abolish its endonuclease activity *in vitro* (2, 3) and DSB generation activity *in vivo* (9).

We then combined the obtained plasmids and strains, induced CbAgo expression, and recovered the induced cells to measure recombination frequencies, which were calculated as the fraction of ampicillin-resistant cells that became resistant to kanamycin (kanamycin-resistant and ampicillin-resistant colony-forming units (cfu)/ampicillin-resistant cfu), because only the recombinants have restored functional kanamycin resistance gene. When there are three or six Chi sites adjacent to the recombination cassette (corresponding strain 3×ChikanS and 6×ChikanS), recombination frequencies by CbAgo/GS are significantly higher than the rest control groups (Supplementary Figure S2A), suggesting a recombination pathway that is mediated by guide-directed CbAgo cleavage.

Interestingly, we observed remarkable recombination frequencies in dCbAgo/GS groups in some conditions (Supplementary Figure S2A). When there is no Chi site adjacent to the recombination cassette (corresponding strain nonChikanS), the cell bearing dCbAgo/GS had a recombination frequency being ∼10-fold higher than the one bearing CbAgo/GS. These results suggest there is a recombination pathway that is mediated by the non-cleavage function of CbAgo and is outperformed by the cleavage-dependent pathway in the presence of the CbAgo active site. Although the exact mechanism remains unknown, this dCbAgo-mediated recombination pathway should be independent of DSB and RecBCD because dCbAgo/GS was previously shown not able to generate DSB or trigger RecBCD activity *in vivo* (9).

We also examined the effects of GS length on the recombination frequency (Supplementary Figure S2B) and found that in the range of 50 to 500 bp, recombination frequency increases as GS length increases. We decided to use 1000 bp as the GS length and 3×ChikanS as the model strain to perform fluctuation analysis (25) to estimate recombination rates (Figure 1D). The cell bearing CbAgo/GS had a recombination rate that is 5-fold higher than the ones with dCbAgo, with and without GS, 15-fold higher than the one with CbAgo-only, and 30-fold higher over the rest control groups. The actual contribution of dCbAgo-mediated recombination to the total recombination events in 3×ChikanS_CbAgo/GS should be much smaller than one-fifth because it should be largely outperformed by the cleavage-dependent pathway as previous observation suggests. Together, these findings reveal novel chromosome recombination that is induced by guide-directed CbAgo cleavage. Its Chi site dependence implies DSB formation and RecBCD processing during recombination.

### Validation of the recombination system

To demonstrate the reliability of our recombination system, we tested four additional pAgos in strain 3×ChikanS, including CaAgo, CdAgo, CpAgo, and IbAgo (refer to Supplementary Table S4 for the summary of pAgos used in this study). In the presence of GS, the rates of recombination induced by different pAgos correlate with the rank order of their reported *in vitro* ssDNA cleavage activity (Figure 2A, see ref. 26). We also tested an engineered DSB in our system by integrating a pal246 into the lacZ locus on the genome of strain 3×ChikanS and its *sbcCD* knockout mutant and measuring their recombination rates (Figure 2B, Supplementary Figure S3). The sbcCD^+^, lacZ::pal246 strain yielded a ∼100-fold increase in recombination rate compared to the sbcCD^+^, lacZ^+^ strain and *sbcCD*, lacZ::pal246 strain. The bigger fold change stimulated by SbcCD/pal246 over CbAgo/GS is consistent with the previous observation that SbcCD/pal246 is more efficient in DSB generation than CbAgo/GS *in vivo* (9). These findings indicate a strong correlation between pAgo DNA cleavage activity, DSB generation efficiency, and recombination rate in our system.

**Figure 2.**
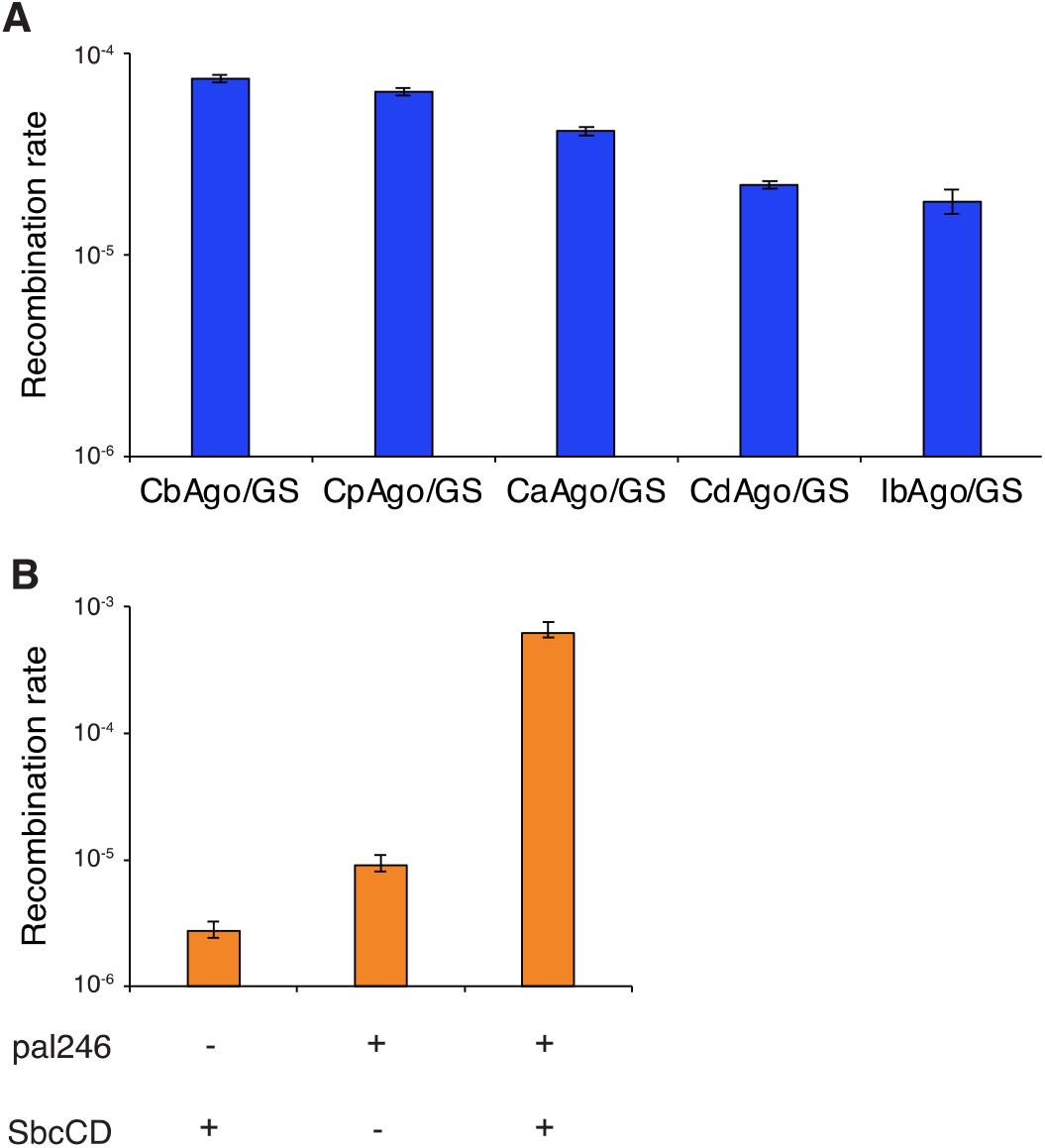
Recombination rate by pAgo/GS correlates with pAgo *in vitro* DNA cleavage activity and *in vivo* DSB generation efficiency. (A) Recombination rates using different pAgos in the presence of GS in strain 3×ChikanS. See ref. 26 for comparisons of *in vitro* DNA cleavage activity among different pAgos. (B) Recombination rates using sbcCD^+^/ΔsbcCD, lacZ^+^/lacZ::pal246 strains. Strains used here were 3×ChikanS, 3×ChikanS_pal246_ΔsbcCD and 3×ChikanS_pal246. See ref. 9 for the comparison of *in vivo* DSB generation efficiency between SbcCD/pal246 and CbAgo/GS. See Supplementary Figure S3 for the genetic structure of engineered DSB. Recombination rates were determined by fluctuation analysis from eight independent cultures. Error bars represent 95% confidence intervals.

### Recombination depends on DSB generation and RecBCD

To gain more insight into our recombination system, we first sought to provide solid evidence that suggests CbAgo can be directed to attack *E. coli* chromosomes and cause DNA damage (Figure 1A, steps i and ii). Since DNA damage induces the cellular SOS response, we used an *E. coli* strain carrying a chromosomally located gfp gene controlled by an SOS-inducible *sulA* promoter (27) and performed flow cytometry to quantify the single-cell fluorescence level (Figure 3A). We observed a 6-fold increase of fluorescence in cells expressing CbAgo and a 15-fold increase in cells containing CbAgo/GS, while the cells containing dCbAgo/GS exhibited no difference in cellular fluorescence level compared to the cells bearing empty plasmids (Figure 3B). The fluorescence increase in cells expressing CbAgo without GS can be explained by previous observations that CbAgo actively degrades plasmids (9) and plasmids degradation triggers SOS-response (28). Importantly, the significant increase of fluorescence in cells expressing CbAgo in the presence of GS confirms that CbAgo can be guided to attack chromosomes, while dCbAgo/GS cannot.

**Figure 3.**
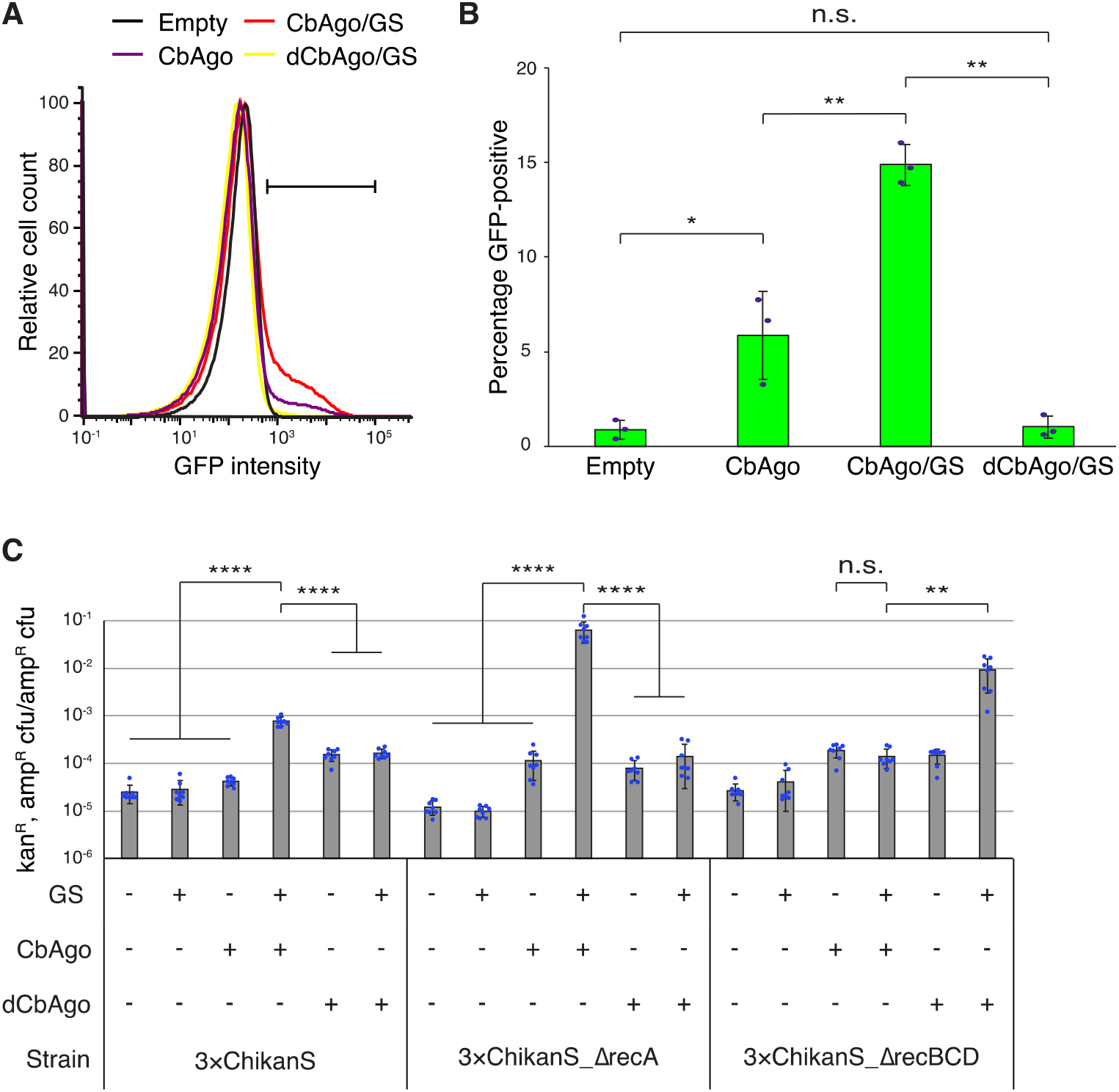
Recombination by CbAgo/GS depends on DSB generation and RecBCD processing but is independent of RecA. (A) CbAgo/GS induces cellular SOS response. Flow cytometry histograms normalized to the mode of the population, combining data from three independent cultures. Horizontal bar represents the GFP^+^ gate. (B) Quantification of GFP-positive cells within the GFP^+^ gate. Error bars, mean ± s.d. from three independent cultures. (C) Recombination frequencies in different genetic contexts. Error bars, mean ± s.d. from eight independent cultures. *P* values were calculated by two-tailed unpaired Student’s t-test; n.s. *P* > 0.05, \**P* < 0.05, \*\**P* < 0.01, \*\*\**P* < 0.001, \*\*\*\**P* < 0.0001.

Then we sought to determine the involvement of *E. coli* endogenous DNA repair machinery RecA and RecBCD in the recombination by creating and testing *recA* and *recBCD* knockout mutants of strain 3×ChikanS. Since the viabilities of knockout strains varied a great deal after induction (Supplementary Figure S4), we determined the fluctuation analysis is no longer suitable and decided to directly analyze recombination frequencies. In the *recBCD* strain, recombination frequency by CbAgo/GS had no difference with the CbAgo-only control, representing a profound change from the result using recBCD^+^ strain (Figure 3C), indicating CbAgo/GS induced recombination depends on RecBCD (Figure 1A, step iii). Since RecBCD works closely with DSB (29), this observation also suggests DSB generation in CbAgo/GS induced recombination. An interesting discovery was that the *recBCD* strain bearing dCbAgo/GS showed ∼1000-fold decreased ampicillin-resistant cfu and only ∼20-fold decreased kanamycin-resistant and ampicillin-resistant cfu compared to its RecBCD^+^ counterpart (Supplementary Figure S4). These changes resulted in increased recombination frequency (Figure 3C), suggesting dCbAgo-mediated recombination is RecBCD-independent. Moreover, this growth inhibition was reduced by ∼60-fold in the presence of the CbAgo active site, suggesting it is outperformed or inhibited when the CbAgo active site is present.

The CbAgo/GS induced DSBs can further explain the extremely low viability of *recA* strain bearing CbAgo/GS (Supplementary Figure S4): without the protection of RecA, continuously introduced DSBs trigger extensive DNA degradation by RecBCD, causing an enormous loss of chromosomal DNA and subsequent cell death (30-32). On the other hand, these cells had a high recombination frequency close to 0.1 (Figure 3C), indicating RecA is not essential in the recombination induced by guide-directed CbAgo cleavage even though it actively repairs DSBs generated during the process. These findings prompt us to modify our model that the final step of recombination (Figure 1A, step iv) should be independent of RecA, reminiscent of the RecA-independent, replication arrest-induced deletion (33,34).

### CbAgo cleavage assists recombineering

The observation that the cfu of *recA* strain was reduced by three orders of magnitude when its genome is targeted by CbAgo (Supplementary Figure S4) is very intriguing, as it supports a strategy to leverage CbAgo/GS targeting as a counter-selection to facilitate recombineering (Figure 4A). For comparison, a self-targeting CRISPR-Cas9 system was reported to reduce cfu by three orders of magnitude in *E. coli* (35). Co-expressing the CRISPR-Cas9 system to eliminate unedited cells, Lambda-Red recombineering achieved an increase of efficiency by ∼10^4^ fold and a 65% overall mutation rate. We sought to combine the CbAgo targeting system with Lambda-Red recombineering by introducing CbAgo expression plasmids into strain SIJ488_ΔrecA, which is RecA-deficient and has arabinose inducible Lambda-Red recombineering genes integrated into its genome. We first performed the standard recombineering procedure with a dsDNA donor encoding kanamycin resistance cassette to replace the genomic lacZ gene. The recombineering efficiency was 1.2 × 10^−4^, calculated from the fraction of cells that became kanamycin resistant. Then we performed recombineering in CbAgo-plasmids contained cells, induced CbAgo expression, and recovered the cells to characterize the proposed counter-selection effect. The cell transformed with pTet_CbAgo/GS had a mutation efficiency of 2.3 × 10^−2^, representing a ∼100-fold increase in efficiency from standard recombineering (Figure 4B). Other control groups did not yield improvement, therefore the increased proportion of the edited cell population depends on guide-directed CbAgo cleavage.

**Figure 4.**
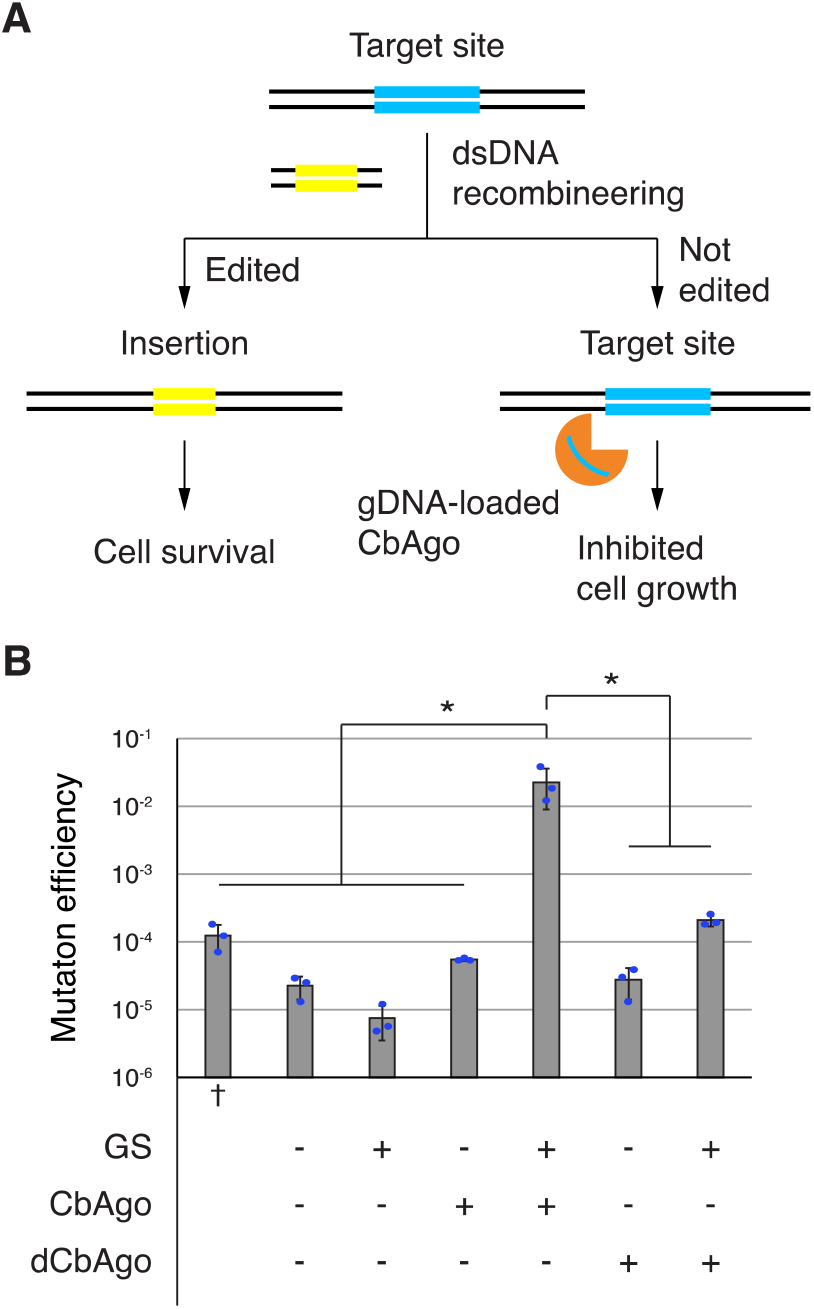
Targeted CbAgo cleavage assists Lambda Red-mediated recombineering in RecA-deficient *E. coli* strain. (A) Scheme of CbAgo-assisted recombineering. After recombineering, the growth of unedited cells will be suppressed by CbAgo cleavage and subsequent DNA degradation, while successfully edited cells will be resistant to CbAgo cleavage and exhibit kanamycin resistance. Yellow, kanamycin resistance cassette. Blue, lacZ gene. (B) Mutation rates using strain SIJ488_ΔrecA in different genetic contexts. †Standard recombineering procedure was applied using plasmid-free cells. See materials and methods for details. Error bars, mean ± s.d. from three independent experiments. *P* values were calculated by two-tailed unpaired Student’s t-test; \**P* < 0.05.

## DISCUSSION

Our study here demonstrates the combination of CbAgo and plasmid-encoded GS can induce mutations in the *E. coli* chromosome, via guide-directed CbAgo cleavage of target DNA, activation of DNA repair mechanism, and subsequent chromosome recombination. This strategy may apply to other organisms if chromosomal DSBs can be introduced and necessary cellular repair machinery can be triggered. Besides, we also demonstrate the potential of pAgo targeting to assist recombineering in RecA-deficient strains as another genome editing strategy. This method may extend to RecA-active strains, if RecA activity can be efficiently inhibited by, for example, expressing RecA inhibitor (36). A recent study also reported NgAgo-assisted recombineering (NgAgo, pAgo from *Natronobacterium gregoryi*), but the fold change was smaller than 2 and the enhancement of editing was not dependent on NgAgo endonuclease activity (37).

The mechanism of guide-directed recombination by dCbAgo in our system remains unknown, although this pathway appears to be independent of RecBCD and DSB. Since dCbAgo can load gDNAs from plasmids *in vivo* (9) (Supplementary Figure. S1), there is a possibility that dCbAgo may play a role in target recognition and following recruitment of *E. coli* nuclease or recombinase. This speculation is supported by findings in other pAgo research (15, 38, 39) and the fact that many pAgo genes have been found associated with a variety of genes including nuclease and helicase (10, 14).

Although CbAgo is among the most active pAgo nucleases identified so far (26), its DNA cleavage activity is still ∼10-fold lower than restriction endonucleases (3). Our recombination system presented here can potentially serve as a reporter and selection platform, which links pAgo cleavage to the development of antibiotic resistance. Therefore, mutants with enhanced DNA cleavage activity are likely to exhibit higher survival rates and be selected. Highly active pAgo nuclease, once obtained, should have bigger potential in DNA-guided genome engineering *in vivo*, and alternatively, may serve as a versatile restriction enzyme *in vitro* with the capability of targeting theoretically any DNA sequence using small oligos as gDNAs, providing a unique advantage over commercial ones (40).

## DATA AVAILABILITY

Additional notes and data are available in the Supplemental materials.

## SUPPLEMENTARY DATA

Supplementary Data are available at NAR online.

## ACKNOWLEDGEMENT

We thank Dr. Alexei A. Aravin, Dr. Andrey Kulbachinskiy, Dr. Daria Esyunina, and Dr. David R. F. Leach for helpful discussions. We thank Rochelle A. Diamond and Jamie Tijerina for their help in flow cytometry experiments.

## Author contributions

S.H. designed and carried out experiments, with K.W. and S.L.M. providing guidance. S.H., K.W., and S.L.M. wrote the manuscript.

## REFERENCES

1. Swarts, D.C., Jore, M.M., Westra, E.R., Zhu, Y., Janssen, J.H., Snijders, A.P., Wang, Y., Patel, D.J., Berenguer, J., Brouns, S.J.J., et al. (2014) DNA-guided DNA interference by a prokaryotic Argonaute. Nature, 507, 258–261.

2. Hegge, J.W., Swarts, D.C., Chandradoss, S.D., Cui, T.J., Kneppers, J., Jinek, M., Joo, C. and van der Oost, J. (2019) DNA-guided DNA cleavage at moderate temperatures by Clostridium butyricum Argonaute. Nucleic Acids Research, 47, 5809–5821.

3. Kuzmenko, A., Yudin, D., Ryazansky, S., Kulbachinskiy, A. and Aravin, A.A. (2019) Programmable DNA cleavage by Ago nucleases from mesophilic bacteria Clostridium butyricum and Limnothrix rosea. Nucleic Acids Research, 47, 5822–5836.

4. Cao, Y., Sun, W., Wang, J., Sheng, G., Xiang, G., Zhang, T., Shi, W., Li, C., Wang, Y., Zhao, F., et al. (2019) Argonaute proteins from human gastrointestinal bacteria catalyze DNA-guided cleavage of single- and double-stranded DNA at 37 °C. Cell Discov, 5, 1–4.

5. Willkomm, S., Oellig, C.A., Zander, A., Restle, T., Keegan, R., Grohmann, D. and Schneider, S. (2017) Structural and mechanistic insights into an archaeal DNA-guided Argonaute protein. Nat Microbiol, 2, 1–7.

6. Sheng, G., Zhao, H., Wang, J., Rao, Y., Tian, W., Swarts, D.C., Oost, J. van der, Patel, D.J. and Wang, Y. (2014) Structure-based cleavage mechanism of Thermus thermophilus Argonaute DNA guide strand-mediated DNA target cleavage. PNAS, 111, 652–657.

7. Liu, Y., Li, W., Jiang, X., Wang, Y., Zhang, Z., Liu, Q., He, R., Chen, Q., Yang, J., Wang, L., et al. (2021) A programmable omnipotent Argonaute nuclease from mesophilic bacteria Kurthia massiliensis. Nucleic Acids Research, 49, 1597–1608.

8. Kropocheva, E., Kuzmenko, A., Aravin, A.A., Esyunina, D. and Kulbachinskiy, A. (2021) A programmable pAgo nuclease with universal guide and target specificity from the mesophilic bacterium Kurthia massiliensis. Nucleic Acids Research, 49, 4054–4065.

9. Kuzmenko, A., Oguienko, A., Esyunina, D., Yudin, D., Petrova, M., Kudinova, A., Maslova, O., Ninova, M., Ryazansky, S., Leach, D., et al. (2020) DNA targeting and interference by a bacterial Argonaute nuclease. Nature, 587, 632–637.

10. Swarts, D.C., Makarova, K., Wang, Y., Nakanishi, K., Ketting, R.F., Koonin, E.V., Patel, D.J. and van der Oost, J. (2014) The evolutionary journey of Argonaute proteins. Nat Struct Mol Biol, 21, 743–753.

11. Ryazansky, S., Kulbachinskiy, A. and Aravin, A.A. The Expanded Universe of Prokaryotic Argonaute Proteins. mBio, 9, e01935–18.

12. Meister, G. (2013) Argonaute proteins: functional insights and emerging roles. Nat Rev Genet, 14, 447–459.

13. Gebert, D. and Rosenkranz, D. (2015) RNA-based regulation of transposon expression. WIREs RNA, 6, 687–708.

14. Makarova, K.S., Wolf, Y.I., van der Oost, J. and Koonin, E.V. (2009) Prokaryotic homologs of Argonaute proteins are predicted to function as key components of a novel system of defense against mobile genetic elements. Biology Direct, 4, 29.

15. Olovnikov, I., Chan, K., Sachidanandam, R., Newman, D.K. and Aravin, A.A. (2013) Bacterial Argonaute Samples the Transcriptome to Identify Foreign DNA. Molecular Cell, 51, 594–605.

16. Swarts, D.C., Hegge, J.W., Hinojo, I., Shiimori, M., Ellis, M.A., Dumrongkulraksa, J., Terns, R.M., Terns, M.P. and van der Oost, J. (2015) Argonaute of the archaeon Pyrococcus furiosus is a DNA-guided nuclease that targets cognate DNA. Nucleic Acids Research, 43, 5120–5129.

17. Swarts, D.C., Koehorst, J.J., Westra, E.R., Schaap, P.J. and Oost, J. van der (2015) Effects of Argonaute on Gene Expression in Thermus thermophilus. PLOS ONE, 10, e0124880.

18. Swarts, D.C., Szczepaniak, M., Sheng, G., Chandradoss, S.D., Zhu, Y., Timmers, E.M., Zhang, Y., Zhao, H., Lou, J., Wang, Y., et al. (2017) Autonomous Generation and Loading of DNA Guides by Bacterial Argonaute. Molecular Cell, 65, 985-998.e6.

19. Hegge, J.W., Swarts, D.C. and van der Oost, J. (2018) Prokaryotic Argonaute proteins: novel genome-editing tools? Nat Rev Microbiol, 16, 5–11.

20. Eykelenboom, J.K., Blackwood, J.K., Okely, E. and Leach, D.R.F. (2008) SbcCD Causes a Double-Strand Break at a DNA Palindrome in the Escherichia coli Chromosome. Molecular Cell, 29, 644–651.

21. Dillingham, M.S. and Kowalczykowski, S.C. (2008) RecBCD Enzyme and the Repair of Double-Stranded DNA Breaks. Microbiology and Molecular Biology Reviews, 72, 642–671.

22. Sinha, A.K., Possoz, C. and Leach, D.R.F. (2020) The Roles of Bacterial DNA Double-Strand Break Repair Proteins in Chromosomal DNA Replication. FEMS Microbiology Reviews, 44, 351–368.

23. Smith, G.R. (2012) How RecBCD Enzyme and Chi Promote DNA Break Repair and Recombination: a Molecular Biologist’s View. Microbiology and Molecular Biology Reviews, 76, 217–228.

24. Wigley, D.B. (2013) Bacterial DNA repair: recent insights into the mechanism of RecBCD, AddAB and AdnAB. Nat Rev Microbiol, 11, 9–13.

25. Hall, B.M., Ma, C.-X., Liang, P. and Singh, K.K. (2009) Fluctuation AnaLysis CalculatOR: a web tool for the determination of mutation rate using Luria–Delbrück fluctuation analysis. Bioinformatics, 25, 1564–1565.

26. Vaiskunaite, R., Vainauskas, J., Morris, J.J.L., Potapov, V. and Bitinaite, J. (2022) Programmable cleavage of linear double-stranded DNA by combined action of Argonaute CbAgo from Clostridium butyricum and nuclease deficient RecBC helicase from E. coli. Nucleic Acids Research, 50, 4616–4629.

27. Pennington, J.M. and Rosenberg, S.M. (2007) Spontaneous DNA breakage in single living Escherichia coli cells. Nat Genet, 39, 797–802.

28. Citorik, R.J., Mimee, M. and Lu, T.K. (2014) Sequence-specific antimicrobials using efficiently delivered RNA-guided nucleases. Nat Biotechnol, 32, 1141–1145.

29. Taylor, A.F. and Smith, G.R. (1985) Substrate specificity of the DNA unwinding activity of the RecBC enzyme of Escherichia coli. Journal of Molecular Biology, 185, 431–443.

30. Capaldo, F.N. and Barbour, S.D. (1975) DNA content, synthesis and integrity in dividing and non-dividing cells of rec− strains of Escherichia coli K12. Journal of Molecular Biology, 91, 53–66.

31. Kuzminov, A. and Stahl, F.W. (1997) Stability of linear DNA in recA mutant Escherichia coli cells reflects ongoing chromosomal DNA degradation. Journal of Bacteriology, 179, 880–888.

32. Skarstad, K. and Boye, E. (1993) Degradation of individual chromosomes in recA mutants of Escherichia coli. Journal of Bacteriology, 175, 5505–5509.

33. Bierne, H., Ehrlich, S.D. and Michel, B. (1997) Deletions at stalled replication forks occur by two different pathways. The EMBO Journal, 16, 3332–3340.

34. Michel, B. (2000) Replication fork arrest and DNA recombination. Trends in Biochemical Sciences, 25, 173–178.

35. Jiang, W., Bikard, D., Cox, D., Zhang, F. and Marraffini, L.A. (2013) RNA-guided editing of bacterial genomes using CRISPR-Cas systems. Nat Biotechnol, 31, 233–239.

36. Moreb, E.A., Hoover, B., Yaseen, A., Valyasevi, N., Roecker, Z., Menacho-Melgar, R. and Lynch, M.D. (2017) Managing the SOS Response for Enhanced CRISPR-Cas-Based Recombineering in E. coli through Transient Inhibition of Host RecA Activity. ACS Synth. Biol., 6, 2209–2218.

37. Lee, K.Z., Mechikoff, M.A., Kikla, A., Liu, A., Pandolfi, P., Fitzgerald, K., Gimble, F.S. and Solomon, K.V. (2021) NgAgo possesses guided DNA nicking activity. Nucleic Acids Research, 49, 9926–9937.

38. Fu, L., Xie, C., Jin, Z., Tu, Z., Han, L., Jin, M., Xiang, Y. and Zhang, A. (2019) The prokaryotic Argonaute proteins enhance homology sequence-directed recombination in bacteria. Nucleic Acids Research, 47, 3568–3579.

39. Jolly, S.M., Gainetdinov, I., Jouravleva, K., Zhang, H., Strittmatter, L., Bailey, S.M., Hendricks, G.M., Dhabaria, A., Ueberheide, B. and Zamore, P.D. (2020) Thermus thermophilus Argonaute Functions in the Completion of DNA Replication. Cell, 182, 1545-1559.e18.

40. Enghiad, B. and Zhao, H. (2017) Programmable DNA-Guided Artificial Restriction Enzymes. ACS Synth. Biol., 6, 752–757.

