## Supplementary Information for "Genome manipulation by guide-directed Argonaute cleavage"

**DNA sequence of the recombination cassette.**

Cyan: EM7 promoter, Yellow: kanamycin direct repeat sequence, Red: stop codon array, Green: kanamycin C-terminus sequence

CCACTAGTTAGCTCGAGGGTGTGGAAAGTCCCCAGGCTCCCCAGCAGGCAGAAGTATGCAAAG
CATGCATCTCAATTAGTCAGCAACCAGGTGTGGAAAGTCCCCAGGCTCCCCAGCAGGCAGAAGT
ATGCAAAGCATGCATCTCAATTAGTCAGCAACCATAGTCCCGCCCCTAACTCCGCCCATCCCGC
CCCTAACTCCGCCCAGTTCCGCCCATTCTCCGCCCCATGGCTGACTAATTTTTTTTATTTATGCAG
AGGCCGAGGCCGCCTCTGCCTCTGAGCTATTCCAGAAGTAGTGAGGAGGCTTTTTTGGAGGCCT
AGGCTTTTGCAAAAAGCTCCCGGGAGCTTGTATATCCATTTTCGGATCTGATCAGCACGTGTTGA
CAATTAATCATCGGCATAGTATATCGGCATAGTATAATACGACAAGGTGAGGAACTAAACCATGA
GCCATATTCAACGGGAAACGTCTTGCTCTAGGCCGCGATTAAATTCCAACATGGATGCTGATTTA
TATGGGTATAAATGGGCTCGCGATAATGTCGGGCAATCAGGTGCGACAATCTATCGATTGTATGG
GAAGCCCGATGCGCCAGAGTTGTTTCTGAAACATGGCAAAGGTAGCGTTGCCAATGATGTTACA
GATGAGATGGTCAGACTAAACTGGCTGACGGAATTTATGCCTCTTCCGACCATCAAGCATTTTAT
CCGTACTCCTGATTAACTGACTAGCATATTCAACGGGAAACGTCTTGCTCTAGGCCGCGATTAAA
TTCCAACATGGATGCTGATTTATATGGGTATAAATGGGCTCGCGATAATGTCGGGCAATCAGGTG
CGACAATCTATCGATTGTATGGGAAGCCCGATGCGCCAGAGTTGTTTCTGAAACATGGCAAAGG
TAGCGTTGCCAATGATGTTACAGATGAGATGGTCAGACTAAACTGGCTGACGGAATTTATGCCTC
TTCCGACCATCAAGCATTTTATCCGTACTCCTGATGATGCATGGTTACTCACCACTGCGATCCCC
GGGAAAACAGCATTCCAGGTATTAGAAGAATATCCTGATTCAGGTGAAAATATTGTTGATGCGCT
GGCAGTGTTCCTGCGCCGGTTGCATTCGATTCCTGTTTGTAATTGTCCTTTTAACAGCGATCGCG
TATTTCGTCTCGCTCAGGCGCAATCACGAATGAATAACGGTTTGGTTGATGCGAGTGATTTTGAT
GACGAGCGTAATGGCTGGCCTGTTGAACAAGTCTGGAAAGAAATGCATAAACTTTTGCCATTCTC
ACCGGATTCAGTCGTCACTCATGGTGATTTCTCACTTGATAACCTTATTTTTGACGAGGGGAAATT
AATAGGTTGTATTGATGTTGGACGAGTCGGAATCGCAGACCGATACCAGGATCTTGCCATCCTAT
GGAACTGCCTCGGTGAGTTTTCTCCTTCATTACAGAAACGGCTTTTTCAAAAATATGGTATTGATA
ATCCTGATATGAATAAATTGCAGTTTCATTTGATGCTCGATGAGTTTTTCTAACACGTGCTACGAG
ATTTCGATTCCACCGCCGCCTTCTATGAAAGGTTGGGCTTCGGAATCGTTTTCCGGGACGCCGG
CTGGATGATCCTCCAGCGCGGGGATCTCATGCTGGAGTTCTTCGCCCACCCCAACTTGTTTATT
GCAGCTTATAATGGTTACAAATAAAGCAATAGCATCACAAATTTCACAAATAAAGCATTTTTTTCAC
TGCATTCTAGTTGTGGTTTGTCCAAACTCATCAATGTATCTTATCATGTCTGAATTCCCGGGGATC

**DNA sequence of the guide sequence.**

CACTGGCCGTCGTTTTACAACGTCGTGACTGGGAAAACCCTGGCGTTACCCAACTTAATCGCCTT
GCAGCACATCCCCCTTTCGCCAGCTGGCGTAATAGCGAAGAGGCCCGCACCGATCGCCCTTCC
CAACAGTTGCGCAGCCTGAATGGCGAATGGCGCTTTGCCTGGTTTCCGGCACCAGAAGCGGTG
CCGGAAAGCTGGCTGGAGTGCGATCTTCCTGAGGCCGATACTGTCGTCGTCCCCTCAAACTGGC
AGATGCACGGTTACGATGCGCCCATCTACACCAACGTGACCTATCCCATTACGGTCAATCCGCC
GTTTGTTCCCACGGAGAATCCGACGGGTTGTTACTCGCTCACATTTAATGTTGATGAAAGCTGGC
TACAGGAAGGCCAGACGCGAATTATTTTTGATGGCGTTAACTCGGCGTTTCATCTGTGGTGCAAC
GGGCGCTGGGTCGGTTACGGCCAGGACAGTCGTTTGCCGTCTGAATTTGACCTGAGCGCATTTT
TACGCGCCGGAGAAAACCGCCTCGCGGTGATGGTGCTGCGCTGGAGTGACGGCAGTTATCTGG
AAGATCAGGATATGTGGCGGATGAGCGGCATTTTCCGTGACGTCTCGTTGCTGCATAAACCGAC
TACACAAATCAGCGATTTCCATGTTGCCACTCGCTTTAATGATGATTTCAGCCGCGCTGTACTGG
AGGCTGAAGTTCAGATGTGCGGCGAGTTGCGTGACTACCTACGGGTAACAGTTTCTTTATGGCA
GGGTGAAACGCAGGTCGCCAGCGGCACCGCGCCTTTCGGCGGTGAAATTATCGATGAGCGTGG
TGGTTATGCCGATCGCGTCACACTACGTCTGAACGTCGAAAACCCGAAACTGTGGAGCGCCGAA

ATCCCGAATCTCTATCGTGCGGTGGTTGAACTGCACACCGCCGACGGCACGCTGATTGAAGCAG
AAGCCTGCGATGTCGGTTTCCGCGAGGTGCGGATTGAAAA

**Plasmid construction**

pTet_wtCas9 was a gift from Dr. Stanley Qi (Addgene plasmid #44250). pET28b_CbAgo and pET28b_dCbAgo were gifts from Dr. Alexei Aravin. pDL1999 was a gift from Dr. David Leach.

Plasmids pnonChikanS, pChikanS, p3×ChikanS, and p6×ChikanS were created to serve as templates for PCR amplification to generate dsDNA donors for Lambda-Red recombineering.

Plasmid pnonChikanS was constructed as follows. Plasmid backbone consisting of two homology arms to the cynX gene and an EM7 promoter was amplified from pDL1999 using primers kan-bb.F/kan-bb.R. Two copies of kan gene were amplified from plasmid pET28b using primers kan1.F/kan1.R and kan2.F/kan2.R, respectively. Each PCR product was gel-purified and all three fragments were fused together by Gibson Assembly. In this way, the recombination cassette containing two non-functional copies of the kanamycin resistance gene separated by a stop codon array was created and inserted adjacent to the EM7 promoter, between the two cynX homology arms.

Plasmid pChikanS was obtained through amplification of pnonChikanS with primers Chi.F/Chi.R. PCR product was gel-purified and self-ligated by Gibson assembly.

Plasmid p3×ChikanS was obtained through amplification of pChikanS with primers 3×Chi.F/3×Chi.R. PCR product was gel-purified and self-ligated by Gibson assembly.

Plasmid p6×ChikanS was obtained through amplification of p3×ChikanS with primers 6×Chi.F/6×Chi.R. PCR product was gel-purified and self-ligated by Gibson assembly.

To construct plasmids pTet_CbAgo and pTet_dCbAgo, plasmid backbone was amplified from pTet_wtCas9 using primers pTet-bb.F/pTet-bb.R. CbAgo and dCbAgo sequences were amplified from pET28b_CbAgo and pET28b_dCbAgo respectively using primers Cb.F/Cb.R, gel-purified and ligated with the plasmid backbone individually via Gibson Assembly.

To construct plasmids pTet_CbAgo/GS and pTet_dCbAgo/GS, a 1000-bp sequence homologous to lacZ gene was amplified from DL1777 genomic DNA via colony PCR using primers GS-lacZ.F/GS-lacZ.R. Plasmid backbones of pTet_CbAgo and pTet_dCbAgo were also PCR amplified with primers GS-bb.F/GS-bb.R, gel-purified and ligated with the 1000-bp lacZ homologous sequence individually via Gibson Assembly.

To construct plasmids pTet_CpAgo/GS, pTet_CaAgo/GS, pTet_CdAgo/GS, pTet_IbAgo/GS, plasmid backbones were amplified from pTet_CbAgo/GS using primers pTet-bb.F/pTet-bb.R. CpAgo, CaAgo, CdAgo, IbAgo gene fragments were ordered from IDT (Integrated DNA Technologies) or Twist Bioscience, amplified using primers pTet-Ago.F/pTet-Ago.R individually, gel-purified and ligated with the plasmid backbones individually via Gibson Assembly. To construct plasmids pEmpty and pGS, pTet_CbAgo and pTet_CbAgo/GS were amplified using primers empty.F/empty.R respectively to remove the CbAgo gene from the plasmid. The resulting PCR products were gel-purified and self-ligated via Gibson Assembly.

Plasmid pCas9-Red was purchased from Sigma-Aldrich (Catalog Number CAS9BAC1P), which contains the gene for Cas9 from *Streptococcus pyogenes* (spCas9) expressed from its native promoter, and the genes for Lambda-Red recombinases Exo, Beta, and Gam under control of the arabinose-inducible ParaB promoter.

To construct plasmid pgRNA_cynX, plasmid pgRNA_lacZ was used as the template, which encodes a CRISPR-gRNA targeting lacZ under control of the J23119 promoter and a sacB gene from *Bacillus subtilis* for counter-selection. Plasmid backbone was amplified from pgRNA_lacZ using primers gRNA-cynX.F/gRNA-cynX.R, gel-purified, and self-ligated by Gibson Assembly, resulting in the replacement of the original spacer sequence with the synthetic spacer sequence targeting cynX.

**Strain construction**

Strain DL1777, DL2859, and DL2874 were gifts from Dr. David Leach. SMR6669 was a gift from Dr. Susan Rosenberg. SIJ488 was a gift from Dr. Alex Nielsen (Addgene bacterial strain #68246).

To generate strain nonChikanS, ChikanS, 3×ChikanS, 6×ChikanS, plasmid pnonChikanS, pChikanS, p3×ChikanS, and p6×ChikanS were amplified with primers cynX-arm.F/cynX-arm.R respectively and PCR products were gel-purified as dsDNA donors. DL1777 was transformed with pCas9-Red, plated on LB plates supplemented with kanamycin, and incubated overnight at 30 °C. One of the transformants was inoculated in 5 mL LB media supplemented with kanamycin and grown at 30 °C until OD600 = 0.3-0.4. The Lambda-Red genes were then induced with 15 mM L-arabinose for 45 min. The culture was used to prepare electrocompetent cells by washing twice with 10% glycerol and resuspending in 50 µl 10% glycerol. 5 µl mixture of ~250 ng dsDNA and 100 ng plasmid pgRNA_lacZ was added to the cells, which were then subject to electroporation and allowed to recover in 1 ml LB for 2 h at 30 °C. The recovered cells were plated on LB plates supplemented with ampicillin and kanamycin and incubated overnight at 30 °C. Colonies with correct genomic integration were verified by colony PCR and sequencing. Upon obtaining positive hits, plasmids were cured by growing the cells overnight in LB media without antibiotics at 37 °C and plating on LB plates supplemented with 5% sucrose. Successful plasmid curing was verified by colony PCR.

Strain 3×ChikanR was generated by previously described CbAgo/GS mediated recombination using strain 3×ChikanS. Kanamycin-resistant colonies with correct recombination were verified by colony PCR and sequencing. Plasmids were then cured by growing the cells overnight in LB media without antibiotics at 37 °C and plating on LB plates. Successful plasmid curing was confirmed by the cell sensitivity to ampicillin.

To generate strain 3×ChikanS_pal246 and 3×ChikanS_pal246_ΔsbcCD, genomic DNA of 3×ChikanS was amplified via colony PCR with primers cynX-arm.F/gmR-arm.R and PCR product was gel-purified as dsDNA donor. DL2859 and DL2874 were individually transformed with pCas9-Red, plated on LB plates supplemented with kanamycin, and incubated overnight at 30 °C. One of the transformants was inoculated in 5 mL LB media supplemented with kanamycin and grown at 30 °C until OD600 = 0.3-0.4. The Lambda-Red genes were then induced with 15 mM L-arabinose for 45 min. The culture was used to prepare electrocompetent cells by washing twice with 10% glycerol and resuspending in 50 µl 10% glycerol. 5 µl mixture of ~250 ng dsDNA and 100 ng plasmid pgRNA_lacZ was added to the cells, which were then subject to electroporation and allowed to recover in 1 ml LB for 2 h at 30 °C. The recovered cells were plated on LB plates supplemented with ampicillin and kanamycin and incubated overnight at 30 °C. Colonies with correct genomic integration were verified by colony PCR and sequencing. Upon obtaining positive hits, plasmids were cured by growing the cells overnight in LB media without antibiotics at 37 °C and plating on LB plates supplemented with 5% sucrose. Successful plasmid curing was verified by colony PCR.

To generate strain 3×ChikanS_ΔrecBCD and 3×ChikanS_ΔrecA, chloramphenicol-resistance cassettes were amplified from plasmid pDL1999 using primers recA.cmR.F/recA.cmR.R and recBCD.cmR.F/recBCD.cmR.R respectively. PCR products were gel-purified and used as dsDNA donors. 3×ChikanS was transformed with pKD46 plasmid, plated on LB plates supplemented with ampicillin, and incubated overnight at 30 °C. A 5 ml culture inoculated from single colony was grown at 30°C until OD600 = 0.3-0.4. The Lambda-Red genes were then induced with 15 mM L-arabinose for 45 min. The culture was used to prepare electrocompetent cells by washing twice with 10% glycerol and resuspending in 50 µl 10% glycerol. These cells were transformed with ~250 ng dsDNA by electroporation and allowed to recover in 1 mL LB for 2 h at 30 °C. The recovered cells were plated on LB plates supplemented with chloramphenicol and incubated overnight at 37 °C. The correct knockouts were verified by colony PCR and sequencing. Upon obtaining positive hits, the pKD46 plasmid was cured by growing the cell in LB media at 37°C overnight and plating on LB plates. Successful plasmid curing was confirmed by the cell sensitivity to ampicillin.

To generate strain SIJ488_ΔrecA, the chloramphenicol-resistance cassette was amplified from plasmid pDL1999 using primers recA.cmR.F/recA.cmR.R. PCR products were gel-purified and used as dsDNA donor. A 5 ml culture inoculated from single colony of SIJ488 was grown at 37°C until OD600 = 0.3-0.4. The Lambda-Red genes were then induced with 15 mM L-arabinose for 45 min. The culture was used to prepare electrocompetent cells by washing twice with 10% glycerol and resuspending in 50 µl 10% glycerol. These cells were transformed with ~250 ng dsDNA by electroporation and allowed to recover in 1 mL LB for 2 h at 37 °C. The recovered cells were plated on LB plates supplemented with chloramphenicol and incubated overnight at 37 °C. Correct knockouts were verified by colony PCR and sequencing.

To generate strain SIJ488_ΔlacZ, the kanamycin-resistance cassette was amplified from genomic DNA of 3×ChikanR via colony PCR with primers lacZ.kanR.F/lacZ.kanR.R, and the PCR product was gel-purified as dsDNA donor. A 5 ml culture inoculated from single colony of SIJ488 was grown at 37°C until OD600 = 0.3-0.4. The Lambda-Red genes were then induced with 15 mM L-arabinose for 45 min. The culture was used to prepare electrocompetent cells by washing twice with 10% glycerol and resuspending in 50 µl 10% glycerol. These cells were transformed with ~250 ng dsDNA by electroporation and allowed to recover in 1 mL LB for 2 h at 37 °C. The recovered cells were plated on LB plates supplemented with kanamycin and incubated overnight at 37 °C. Correct knockouts were verified by colony PCR and sequencing.

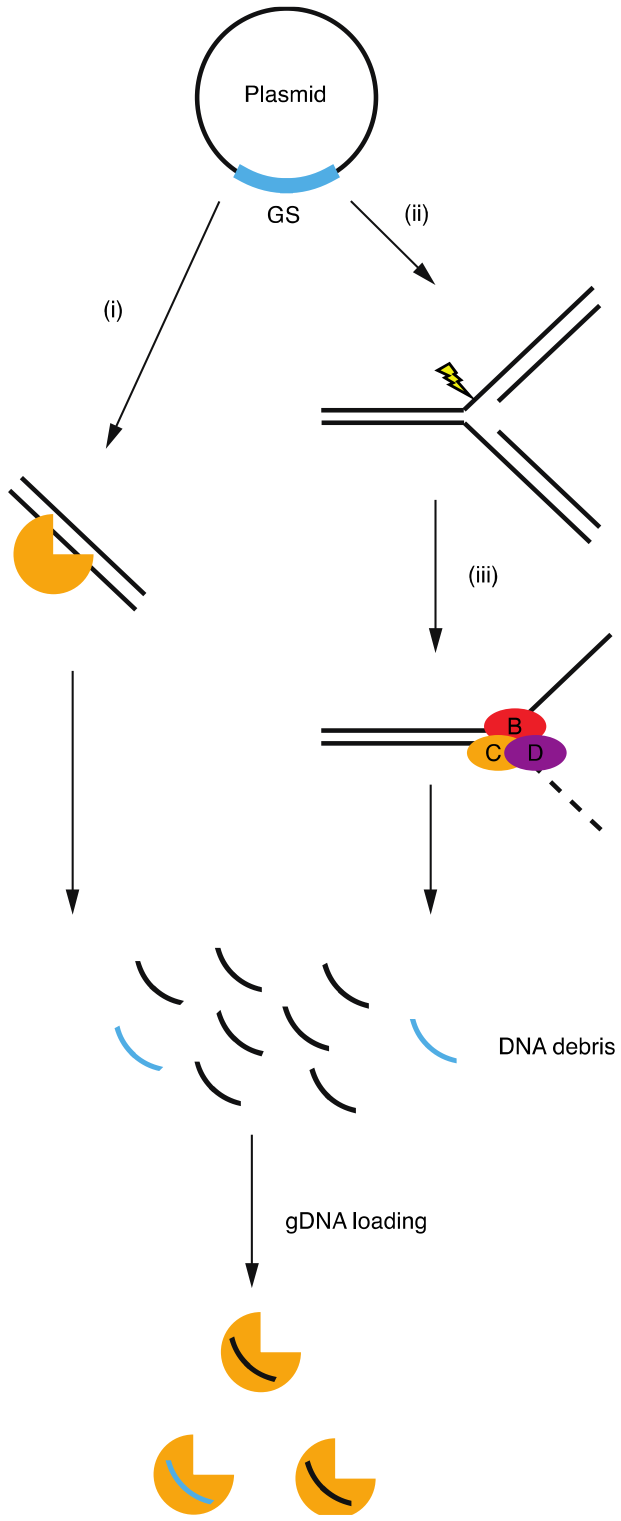


**Figure S1.** Proposed model for gDNA biogenesis of CbAgo. As found in the previous report, CbAgo generates and binds small gDNA from plasmids (1). Small DNA fragments can be generated from guide-free CbAgo mediated plasmid degradation ("chopping" activity), likely at partially unwound dsDNA in AT-rich region (step i) (2). Alternatively, RecBCD-dependent plasmid degradation during plasmid replication (step ii and iii) can also generate small DNA debris, a process similar to the spacer acquisition in CRISPR adaptation (3-5). The resulting small DNA fragments can serve as substrates for gDNA loading by CbAgo. The RecBCD-dependent plasmid degradation may also be responsible for dCbAgo gDNA loading.

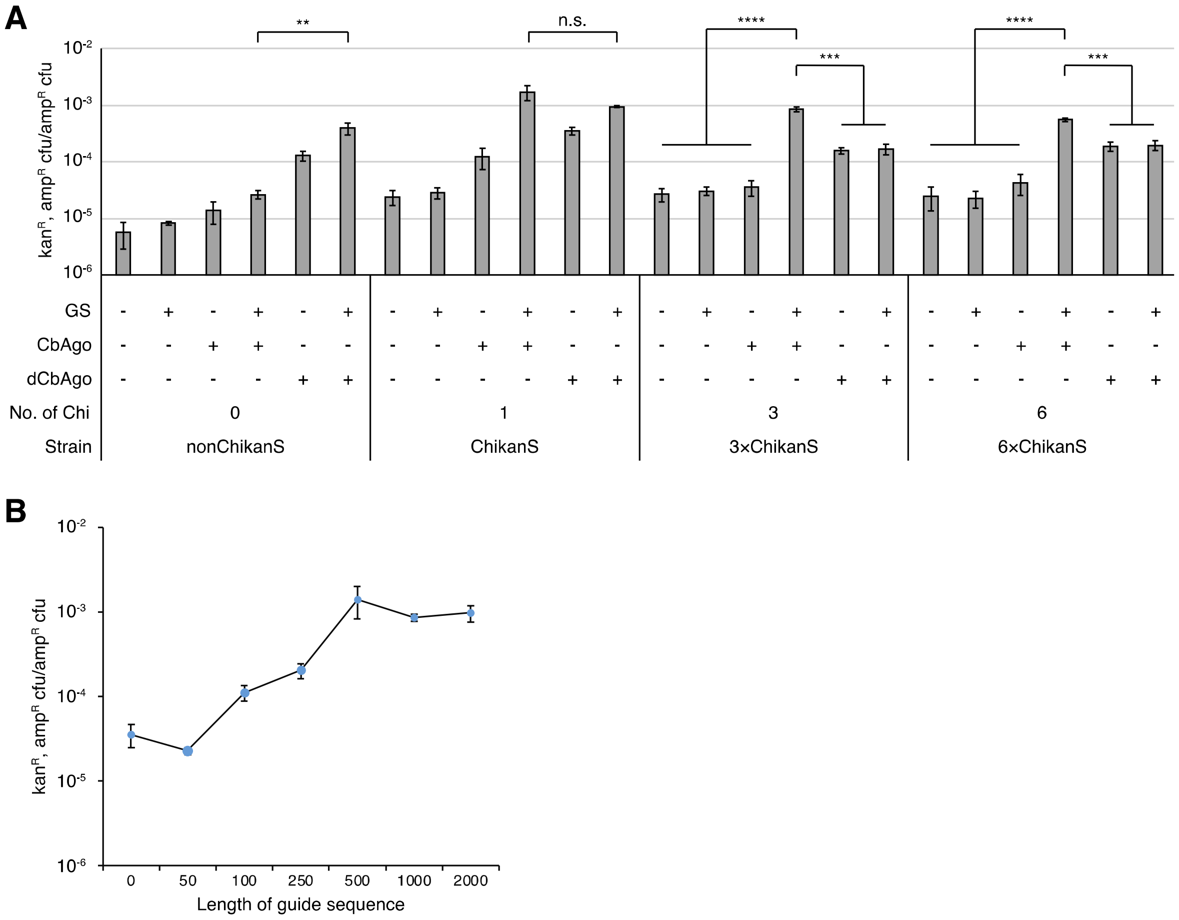


**Figure S2.** Effects of the number of Chi sites and GS length on the recombination frequency. (A) Recombination frequencies in different genetic contexts. Error bars, mean ± s.d. from three independent cultures. (B) Recombination frequencies of strain 3×ChikanS expressing CbAgo in the presence of GS with varying lengths. Error bars, mean ± s.d. from three independent cultures. n.s. P > 0.05, *P < 0.05, **P < 0.01, ***P < 0.001, ****P < 0.0001.

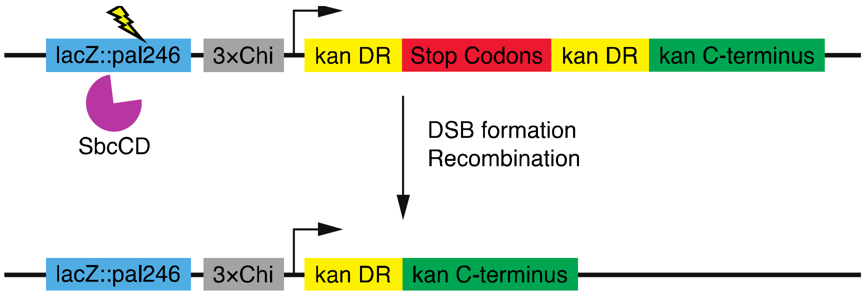


**Figure S3.** Genetic structure of engineered DSB. A pal246 was inserted into lacZ loci, whose cleavage by *E. coli* endogenous nuclease SbcCD forms DSB and stimulates recombination.

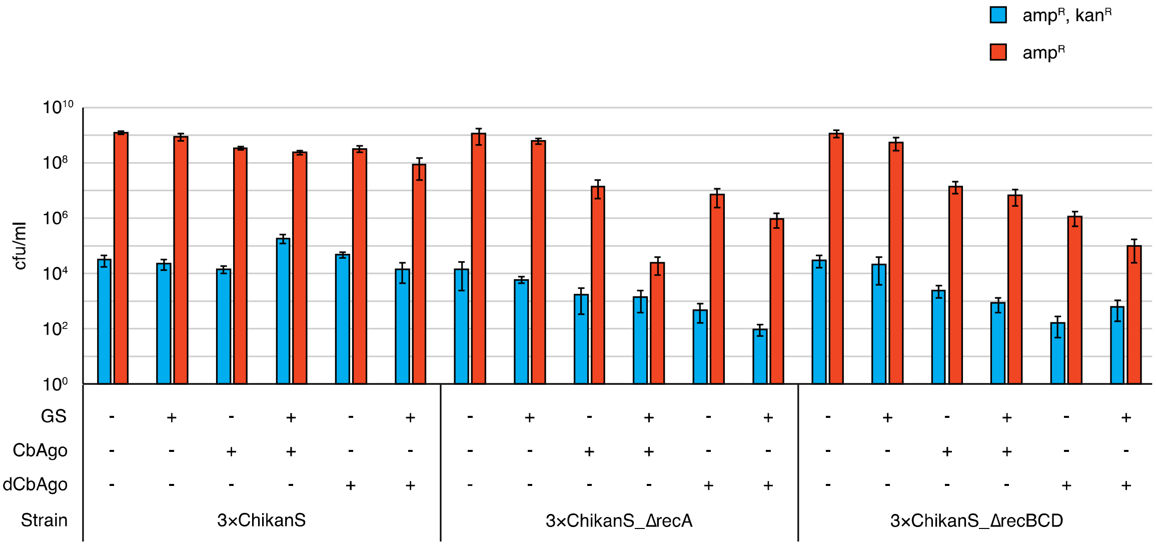


**Figure S4.** Viabilities of cells in different genetic contexts. Error bars, mean ± s.d. from eight independent cultures.

**Table S1.** Strains used in this study.

| Name | Description | Source |
|---|---|---|
| DL1777 | MG1655 *lacIq lacZχ⁻ fnr-267 (∆ynaJ ∆ydaA ∆frn ∆ogt ∆abgT ∆abgB ∆abgA ∆abgR ∆ydaL ∆ydaM ∆dboA ∆ydaO)* | (6) |
| DL2859 | DL1777 *lacZ::pal246 cynX::Gm*$^{R}$ | (6) |
| DL2874 | DL2859 ∆*sbcCD* | (6) |
| nonChikanS | DL1777 *cynX*::[χ⁻ *kan cassette*] | This study |
| ChikanS | DL1777 *cynX*::[χ *kan cassette*] | This study |
| 3×ChikanS | DL1777 *cynX*::[χχχ *kan cassette*] | This study |
| 6×ChikanS | DL1777 *cynX*::[χχχχχχ *kan cassette*] | This study |
| 3×ChikanS_pal246 | 3×ChikanS *lacZ::pal246* | This study |
| 3×ChikanS_pal246_∆sbcCD | 3×ChikanS_pal246 ∆*sbcCD* | This study |
| 3×ChikanS_∆recA | 3×ChikanS ∆*recA::cm*$^{R}$ | This study |
| 3×ChikanS_∆recBCD | 3×ChikanS ∆*recBCD::cm*$^{R}$ | This study |
| 3×ChikanR | DL1777 *cynX::*χχχ *kan*$^{R}$ | This study |
| SMR6669 | MG1655 Δ*att*λ::psulA-*gfpmut2* | (7) |
| SIJ488 | MG1655 Tn7*::*para-exo-beta-gam; prha-FLP; xylSpm-IsceI | Addgene |
| SIJ488_∆recA | SIJ488 ∆*recA::cm*$^{R}$ | This study |
| SIJ488_∆lacZ | SIJ488 ∆*lacZ::kan*$^{R}$ | This study |

**Table S2.** Plasmids used in this study.

| Name | Description | Source |
|---|---|---|
| pET28b | KmR; empty vector | Lab plasmid collection |
| pET28b_CbAgo | KmR; pET28b-derived expression vector with N-terminally 6xHis-tagged codon-optimized CbAgo | A. Aravin Lab |
| pET28b_dCbAgo | KmR; pET28b-derived expression vector with codon-optimized N-terminally 6xHis-tagged catalytically dead CbAgo | A. Aravin Lab |
| pDL1999 | CmR Ts SucS; pTOF24 + *cynX*::[*χ- zeo cassette*] fragment | (6) |
| pnonChikanS | CmR Ts SucS; pTOF24 + *cynX*::[*χ- kan cassette*] fragment | This study |
| pChikanS | CmR Ts SucS; pTOF24 + *cynX*::[*χ kan cassette*] fragment | This study |
| p3×ChikanS | CmR Ts SucS; pTOF24 + *cynX*::[*χχχ kan cassette*] fragment | This study |
| p6×ChikanS | CmR Ts SucS; pTOF24 + *cynX*::[χχχχχχ *kan cassette*] fragment | This study |
| pCas9-Red | KmR Ts; encodes Cas9 from Streptococcus pyogenes (spCas9) expressed from its native promoter, genes for λ-red recombinases exo, beta, and gam under control of the arabinose-inducible ParaB promoter | Sigma Aldrich |
| pgRNA_lacZ | AmpR SucS; encodes spCas9 gRNA targeting lacZ with a spacer sequence 5'-CGGCCAGTGAATCCGTAATCA-3' | Lab plasmid collection |
| pgRNA_cynX | AmpR SucS; encodes spCas9 gRNA targeting cynX with a spacer sequence 5'-CTCTGTGCAACCGGCTATTGC-3' | This study |
| pTet_wtCas9 | AmpR; encodes *S. pyogenes* Cas9 under control of pTet promoter | Addgene |
| pTet_CbAgo | AmpR; encodes CbAgo under control of pTet promoter | This study |
| pTet_dCbAgo | AmpR; encodes dCbAgo under control of pTet promoter | This study |
| pTet_CbAgo/GS | AmpR; encodes CbAgo under control of pTet promoter and contains a 1000 bp sequence homologous to lacZ | This study |
| pTet_dCbAgo/GS | AmpR; encodes dCbAgo under control of pTet promoter and | This study |

| | | |
|---|---|---|
| | contains a 1000 bp sequence homologous to lacZ | |
| pTet_CaAgo/GS | AmpR; encodes CaAgo under control of pTet promoter and contains a 1000 bp sequence homologous to lacZ | This study |
| pTet_CdAgo/GS | AmpR; encodes CdAgo under control of pTet promoter and contains a 1000 bp sequence homologous to lacZ | This study |
| pTet_CpAgo/GS | AmpR; encodes CpAgo under control of pTet promoter and contains a 1000 bp sequence homologous to lacZ | This study |
| pTet_IbAgo/GS | AmpR; encodes IbAgo under control of pTet promoter and contains a 1000 bp sequence homologous to lacZ | This study |
| pGS | AmpR; contains a 1000 bp sequence homologous to lacZ | This study |
| pEmpty | AmpR; empty vector | This study |
| pKD46 | AmpR Ts; encodes genes for Lambda-Red recombinases Exo, Beta, and Gam under control of the arabinose-inducible ParaB promoter | R. Phillips Lab |

**Table S3.** Oligonucleotides used in this study.

| Name | Sequence 5'-3' |
|---|---|
| gRNA-cynX.F | GCAATAGCCGGTTGCACAGAGAGCTAGCATTATACCTAGGA |
| gRNA-cynX.R | TCTGTGCAACCGGCTATTGCGTTTTAGAGCTAGAAATAGCAAG |
| kan1.F | GACAAGGTGAGGAACTAAACCATGAGCCATATTCAACGGG |
| kan1.R | TTGAATATGCTAGTCAGTTAATCAGGAGTACGGATAAAATGC |
| kan2.F | TAACTGACTAGCATATTCAACGGGAAACGTC |
| kan2.R | TCGAAATCTCGTAGCACGTGTTAGAAAAACTCATCGAGCATC |
| kan-bb.F | CACGTGCTACGAGATTTCG |
| kan-bb.R | GGTTTAGTTCCTCACCTTGTC |
| Chi.F | CCCTGCACCACCAGCTAGCTCGAGGGTGTGGAAAGTCCCC |
| Chi.R | AGCTAGCTGGTGGTGCAGGGAATCGGTTTTATCATCGCCGGGC |
| 3×Chi.F | GCGTGTCCACCAGCTCAGCATCGACCACCAGCCAGGCAGAAGTATGCAAAGC |
| 3×Chi.R | TGCTGAGCTGGTGGACACGCGCTGGCTGGTGGTGCAGGGAATCGGTTTTATCATC |
| 6×Chi.F | CGCCCAATGTCCCACCAGCTTGACCAATACCACCAGCCCCTAACTCCGCCCATCC |
| 6×Chi.R | AGCTGGTGGGACATTGGGCGCTGGTGGACTGATGGCGGCTGGTGGTCGATGCTGAG |
| cynX-arm.F | GTTATTGGCGCGGGTAGTATC |
| cynX-arm.R | TATCAAACACTCGCCTGGTG |
| gmR-arm.R | CGTGCGGTCATCACCTTAGATG |
| Cb.F | CTAAAGAGGAGAAAGGATCTATGGGGGGTTCTCATCATCATCATCATCATGG |
| Cb.R | CCTGGAGATCCTTACTCGAGTTATAAGAAGAACAAACGATTGTCTACTACACCC |
| pTet-Ago.F | ATGACGATAAGGATCCGAGC |

| | |
|---|---|
| pTet-Ago.R | CCTGGAGATCCTTACTCGAG |
| pTet-bb.F | CTCGAGTAAGGATCTCCAGG |
| pTet-bb.R | AGATCCTTTCTCCTCTTTAGATC |
| GS-lacZ.F | AGCTCACTCAAAGGCGGTAACACTGGCCGTCGTTTTACAAC |
| GS-lacZ.R | TGATTCTGTGGATAACCGTATTTTCAATCCGCACCTCGCG |
| GS-bb.F | TACGGTTATCCACAGAATCAGGG |
| GS-bb.R | TTACCGCCTTTGAGTGAGCTG |
| empty.F | GGATCTATGTAACTCGAGTAAGGATCTCCAGGCATC |
| empty.R | GAGTTACATAGATCCTTTCTCCTCTTTAGATCTTTTG |
| recA.cmR.F | AAAAAGCAAAAGGGCCGCAGATGCGACCCTTGTGTATCAAACAAGACGATTACGCCCCGCCCTGCCACTC |
| recA.cmR.R | CAACAGAACATATTGACTATCCGGTATTACCCGGCATGACAGGAGTAAAACCGGGAAGCCCTGGGCCAAC |
| recBCD.cmR.F | GTCGGATGCGACATGCGTAACACTCGTACGTCGCATCCGGCAATTACGTTTACGCCCCGCCCTGCCACTC |
| recBCD.cmR.R | GCGAGATGACCCGCCTGCATTGCCCGAATCGTCAGTAGTCAGGAGCCGCTCCGGGAAGCCCTGGGCCAAC |
| lacZ.kanR.F | GATTTCCTTACGCGAAATACGGGCAGACATGGCCTGCCCGGTTATTATTAGAAAAACTCATCGAGCATCAAATG |
| lacZ.kanR.R | GTGTGGAATTGTGAGCGGATAACAATTTCACACAGGAAACAGCTATGACCTGTTGACAATTAATCATCGGC |
| Lambda.Red.F | TAACCACTTCCAGCGCTGAG |
| Lambda.Red.R | TGGCCCTGCACGCGCCGTCG |

**Table S4.** The list of argonaute proteins used in this study.

| Argonaute | GenBank | Organism |
|---|---|---|
| CaAgo | WP_042399050.1 | *Clostridium saudiense* |
| CbAgo | WP_058142162.1 | *Clostridium butyricum* |
| CdAgo | WP_055276084.1 | *Clostridium disporicum* |
| CpAgo | EHP50500.1 | *Clostridium perfringens WAL-14572* |
| IbAgo | WP_055087491.1 | *Intestinibacter bartlettii* |

**SI References**


1. Kuzmenko,A., Oguienko,A., Esyunina,D., Yudin,D., Petrova,M., Kudinova,A., Maslova,O., Ninova,M., Ryazansky,S., Leach,D., *et al.* (2020) DNA targeting and interference by a bacterial Argonaute nuclease. *Nature*, **587**, 632–637.

2. Kuzmenko,A., Yudin,D., Ryazansky,S., Kulbachinskiy,A. and Aravin,A.A. (2019) Programmable DNA cleavage by Ago nucleases from mesophilic bacteria Clostridium butyricum and Limnothrix rosea. *Nucleic Acids Research*, **47**, 5822–5836.

3. Levy,A., Goren,M.G., Yosef,I., Auster,O., Manor,M., Amitai,G., Edgar,R., Qimron,U. and Sorek,R. (2015) CRISPR adaptation biases explain preference for acquisition of foreign DNA. *Nature*, **520**, 505–510.

4. Ivančić-Baće,I., Cass,S.D., Wearne,S.J. and Bolt,E.L. (2015) Different genome stability proteins underpin primed and naïve adaptation in E. coli CRISPR-Cas immunity. *Nucleic Acids Research*, **43**, 10821–10830.

5. Modell,J.W., Jiang,W. and Marraffini,L.A. (2017) CRISPR–Cas systems exploit viral DNA injection to establish and maintain adaptive immunity. *Nature*, **544**, 101–104.

6. Eykelenboom,J.K., Blackwood,J.K., Okely,E. and Leach,D.R.F. (2008) SbcCD Causes a Double-Strand Break at a DNA Palindrome in the Escherichia coli Chromosome. *Molecular Cell*, **29**, 644–651.

7. Pennington,J.M. and Rosenberg,S.M. (2007) Spontaneous DNA breakage in single living Escherichia coli cells. *Nat Genet*, **39**, 797–802.
